# Effect of Antimicrobial Stewardship with Rapid MALDI-TOF Identification and Vitek 2 Antimicrobial Susceptibility Testing on Hospitalization Outcome

**DOI:** 10.1101/581991

**Authors:** Stephen J Cavalieri, Seunghyug Kwon, Renuga Vivekanandan, Sumaya Ased, Cassara Carroll, Jennifer Anthone, David Schmidt, Maddy Baysden, Christopher J Destache

## Abstract

**Introduction:** Rapid organism identification (ID) and antimicrobial susceptibility testing (AST) along with antibiotic stewardship (ASP) are critical to appropriate treatment. We sought to capture time for bacterial culture and initiation of appropriate therapy for patients, from 2017 (without MALDI-TOF/Vitek 2 and ASP) and 2018 (with MALDI-TOF/Vitek 2 and ASP).

**Methods:** Eligible patients admitted to our hospital with a positive sputum, blood, or urine culture. Sequential patients were retrospectively obtained from March 1 to May 31, 2017. Seventy-seven patients from 2017 were compared to 77 patients from 2018. A time-in-motion study was performed to compare time to identification (ID), AST results, and ASP team intervention for the two periods. Data were entered into SPSS (ver 25) for analysis. Results are reported as mean (± SD) or percentage.

**Results:** Time to organism ID was significantly faster in 2018 (2018 24.9 ± 14.4, 2017 33.8 ± 17 h, p=0.001). Time to AST results was also significantly faster for patients in 2018 compared to 2017 (18.2 ± 14 compared to 28.5 ± 14.9 h, p<0.001). ASP team recommended significantly more adjustments to empiric antimicrobial therapy in 2018 (28% of 2018 vs. 2% in 2017, p< 0.001). Length of hospital stay was significantly shorter in 2018 compared to 2017 (2018 10.7 ± 11.1 days and 2017 15.5 ± 18.1 days, p=0.05).

**Conclusions:** Use of MALDI-TOF/Vitek 2 leads to an average 21.5 h faster ID and AST results that can be acted upon by ASP for appropriate antimicrobial recommendations.

---

Bacterial infections in the healthcare associated setting cause significant morbidity and mortality, especially when initial treatment is sub-optimal. This has been clearly demonstrated especially for Gram-negative bloodstream infections (BSIs).(1) Early effective antimicrobial therapy has demonstrated reductions in mortality in patients with sepsis.(2) With antibiotic resistance on the rise, especially in gram-negative bacilli, it is becoming increasingly difficult for clinicians to select the most optimal empiric antibiotic therapy.^(3)^

Laboratory diagnosis of infections is typically dependent on routine culture and phenotypic identification utilizing Gram stain and analysis of biochemical reactions, along with antimicrobial susceptibility testing (AST).(4) Rapid diagnostic tests that allow for earlier identification of pathogens, and in some situations, early identification of resistance patterns, have been shown to reduce time to attaining optimal antibiotic therapy(5-9). Additionally, with the combination of antimicrobial stewardship intervention, these tests have demonstrated a reduced mortality benefit(9, 10). While these tests have enhanced the management of BSIs, the use of MALDI-TOF and Vitek 2 for organism identification and AST along with ASP more generally was of interest to explore.

CHI Health Nebraska is a 14-hospital health system made up of critical access hospitals (5), community-based hospitals (8) and one academic medical center (400 beds) affiliated with Creighton University Health Science Schools (nursing, medicine, pharmacy, physical and occupational therapy). Recent acquisition of a matrix assisted laser desorption/ionization time-of-flight (MALDI-TOF) for organism identification and Vitek 2 for AST in the centralized microbiology laboratory for all CHI Health hospitals has led us to propose this quasi-experimental time-in-motion study design to determine the effect on patient hospitalization outcome. The primary objective was to determine, using a time-in-motion study design, the effect of integrated use of MALDI-TOF/Vitek 2 and ASP for patients with urine, respiratory (based on sputum culture), or BSI at the academic hospital of CHI Health.

## Methods

### Microbiology

For both time periods, routine cultures of urine, sputum, and blood were performed using standard techniques. For blood cultures, the BACTEC system was used. For the 2017 time period, organism ID and AST were performed primarily using the MicroScan microdilution system performed according to manufacturer’s instructions. MALDI-TOF/Vitek 2 (bioMerieux, Inc. St. Louis, MO) became available for general use for organism ID (MALDI-TOF) and AST (Vitek 2) in March 2018 after significant verification procedures were completed. Microbiological workflow was similar to Huang AM, et al.(6) except we are reporting MALDI-TOF and Vitek 2 AST from urine, blood, and sputum cultures after subculture to solid media and incubated overnight.

Additionally, workflow was adjusted to take maximum advantage of the faster turnaround time for ID and AST in that cultures were continuously examined soon after a minimum of 18 hours of incubation after specimen inoculation with growth of potential pathogens processed on the MALDI-TOF and Vitek 2 as soon as possible afterwards. Results were then reported as they became available immediately after they were verified which occurred from 6:30AM to the end of staffing (11PM weekdays and 5PM weekends). Thus, there were two groups in the study; the control group (2017 time period) and the experimental group (2018 time period). Consecutive patients from both time periods (2017 and 2018) were eligible when one culture type was positive (i.e. urine culture positive without blood culture positive) for urine and sputum cultures. BSI could have more than one site with positive isolates.

### Antimicrobial Stewardship

For the 2017 time period (control group), limited ASP was available from staff pharmacists without dedicated ASP personnel. Staff pharmacists were expected to make interventions as part of their normal workflow. In 2018, dedicated ASP personnel were hired providing robust ASP for all CHI Health hospitals (0.5 FTE ID physician and 2.5 FTE ASP pharmacist) and they replaced staff pharmacist’s interventions. All ASP pharmacists monitored patients through TheraDoc and consulted the ID clinician for all antimicrobial recommendations. ASP pharmacists were available during normal work hours (8-4:30) and made recommendations on patients receiving empiric therapy with carbapenems, daptomycin, double coverage, (double anaerobic coverage), all *C. difficile* positive patients, and all *S. aureus* bacteremias. Since both ASP infectious diseases physician and pharmacists made antimicrobial recommendations to primary care physicians (PCPs), other types of recommendations were suggested including ID consult, TEE, and length of therapy as examples. Both years utilized Biofire BCID (Biofire Diagnostics, Salt Lake City, UT) for rapid detection system on all positive blood cultures. However, in 2018 laboratory technicians increased their daily time frame when Biofire was reported for positive blood cultures (7 am-10 pm). All Biofire results were paged to ASP team. The ASP pharmacist kept the Biofire pager (7-5) and then handed it off to the on-call ID fellow overnigh and weekends. In 2017, Biofire results were reported during normal laboratory hours (7 am – 5 pm) and were reviewed by the staff pharmacists as part of an inbasket work queue in EPIC. ASP pharmacists were available five days/week and used TheraDoc (TheraDoc ver 4.7, Hospira, Lake Forest, IL) for clinical decision support software. ASP pharmacists documented their intervention in EPIC as a progress note. Staff pharmacists as part of their normal duties performed weekend coverage. To summarize, in 2017 (control group) rapid diagnostic testing (Biofire) was used for positive blood cultures only and the Microscan instrument was used for urine and sputum results without ASP team intervention. In 2018, Biofire was used for positive blood cultures and called to ASP team and MALDI-TOF/Vitek 2 was used for urine and sputum cultures and not called to the ASP team once positive.

Consent was not required because the standard of care was evaluated for quality assurance purposes. The control group (March-April 2017) identified adult patients (≥ 19 years of age) with blood, urine, or sputum culture obtained and reported while hospitalized. Charlson comorbidity index was used for parity between 2017 and 2018 data(11).

### Exclusion criteria

Patients who were transferred into the hospital from an outside hospital and were empirically treated for a bacterial infection or sepsis, patients transitioned to comfort care or hospice within the first 72 h of antimicrobial therapy, patients who expired prior to identification of the organism, and contaminant cultures and patients empirically placed in isolation for “rule-out” mycobacterial pneumonia.

### Statistical Analysis

Dichotomous data were analyzed using Chi-square or Fisher’s exact test. Continuous, normally distributed data were analyzed by Student t-test. Continuous, non-normally distributed data were analyzed using Mann Whitney U test. Data were collected from EPIC and entered into Statistical Package for the Social Sciences (SPSS, ver. 25, IBM, NY). Data will be presented as percentages or mean ± standard deviation (SD).

## Results

In the control group (2017), a total of 77 patients (27 blood isolates, 25 sputum, and 25 urine isolates) were retrospectively enrolled between March-May 2017. In the MALDI-TOF/Vitek 2 group, a total of 77 patients (28 blood isolates, 24 sputum, and 25 urine isolates) were concurrently enrolled in the same time frame in 2018. Mean (± SD) patient age and Charlson Comorbidity Index (CCI) were not different between the two groups (Table 1). Additionally, patients were admitted to the intensive care unit 31% in 2017 and 32% in 2018, respectively. Microorganisms isolated included 10 (6.5%) that were polymicrobial and the remaining were monomicrobial isolates. The most common single isolate included *E. coli* (30%), methicillin-sensitive *S. aureus* (MSSA) (9%), *S. pneumoniae* (8%), and *P. aeruginosa* (7%). There were no significant differences in the number of isolates in the study from 2017 compared to 2018. The most common sites of infection from patients in both years included urine (69, 45%), lung (55, 38%), and skin (10, 6.5%).

**Table 1.** Demographics of Study Patients

| Variable | 2017 (without ASP + MALDI-TOF/Vitek 2) | 2018 (with ASP + MALDI-TOF/Vitek 2) |
| --- | --- | --- |
| Age (yrs) | 68.2 ± 15.3 | 65.8 ± 18.5 |
| CCI | 3.5 ± 2.2 | 3.6 ± 2.3 |
| Percent female (%) | 56 | 51 |
| Admitted to ICU (%) | 31 | 32 |
| Isolates from: |  |  |
| Blood culture (n) | 27 | 28 |
| Sputum culture (n) | 25 | 24 |
| Urine culture (n) | 25 | 25 |
| Blood cultures: |  |  |
| Candida spp | 0 | 2 |
| *E. coli* | 6 | 8 |
| Enterococcus spp | 3 | 0 |
| Enterobacter spp. | 1 | 0 |
| Klebsiella spp. | 3 | 1 |
| MRSA | 2 | 1 |
| MSSA | 3 | 5 |
| *S. pneumoniae* | 3 | 3 |
| Proteus spp. | 1 | 1 |
| Serratia spp. | 1 | 0 |
| Streptococcus spp. | 1 | 6 |
| VRE | 0 | 1 |
| Polymicrobial isolates | 3 | 0 |
| Sputum cultures: |  |  |
| Acinetobacter spp. | 0 | 1 |
| Citrobacter spp. | 0 | 1 |
| *E. coli* | 0 | 1 |
| Klebsiella spp. | 2 | 2 |
| *H. influenzae* | 1 | 2 |
| *M. catarrhalis* | 0 | 3 |
| MRSA | 1 | 2 |
| Polymicrobial isolates | 3 | 0 |
| Urine cultures: |  |  |
| Aerococcus | 0 | 1 |
| Candida spp. | 1 | 0 |
| *E. cloacae* | 1 | 0 |
| *E. coli* | 17 | 14 |
| Klebsiella spp. | 1 | 3 |
| *S. pneumoniae* | 1 | 0 |
| Proteus spp. | 1 | 6 |
| Pseudomonas spp. | 1 | 0 |
| Stenotrophomonas spp. | 1 | 0 |

CCI = Charlson comorbidity index; ICU = intensive care unit; MRSA = methicillin resistant *S. aureus*; MSSA = methicillin sensitive *S. aureus*

In 2018, with the addition of MALDI-TOF/Vitek 2 results available, ASP team intervened significantly more often for patients’ empiric antimicrobial therapy compared to 2017 (2018 29%, 2017 2%, p < 0.001). The most common intervention was to change empiric therapy (59%). In 2018, the ASP team made empiric antibiotic recommendations. In bacteremic patients, ASP team changed empiric antimicrobial therapy in 19 of 28 (68%). For patients with positive sputum cultures, the ASP team recommended to increase antimicrobial dose in 15 of 24 (62.5%). In patients with positive urine cultures, the majority (84%) did not require a change in empiric therapy. Additionally, time from the AST results to pharmacists making antimicrobial recommendations averaged 0.13 ± 1.5 h compared to 0.24 ± 0.74 h in 2017. However, hospital staff pharmacists made 2 ASP recommendations in 2017 and ASP team made 22 study recommendations in 2018. All but one ASP recommendations were accepted in 2018.

The time-in-motion data collection for culture ID and AST determination demonstrated significant results comparing 2017 without ASP or MALDI-TOF/Vitek 2 and 2018 with ASP and MALDI-TOF/Vitek 2 (Table 2). The time to receive the specimen in the Microbiology Laboratory for both 2017 and 2018 averaged approximately 5 h. Additionally, time from laboratory receipt of specimen until Gram-stain results averaged 15 h in both 2017 and 2018. However, organism identification was significantly faster averaging 10 h sooner and AST results were available 10 h sooner for MALDI-TOF/Vitek 2 patients compared to 2017 without these instruments. Total time for ASP team to make empiric antimicrobial changes averaged 84 h in 2017 and 62.5 h in 2018, a 21.5 h difference.

**Table 2.** Time Parameters to Identify Organism and Obtain AST

| Time Variable | 2017 (without ASP +<br>MALDI-TOF/Vitek 2) | 2018 (with ASP +<br>MALDI-TOF/Vitek 2) | Statistical<br>significance |
| --- | --- | --- | --- |
| Culture transport<br>time (h) | 5.9 ± 6 | 4.7 ± 4.8 | P > 0.05 |
| Gram stain report<br>time (h) | 15.9 ± 14.2 | 14.9 ± 10.4 | P > 0.05 |
| Identify and report<br>organism (h) | 33.8 ± 17 | 24.9 ± 14.4 | P = 0.001 |
| Perform and report<br>AST (h) | 28.5 ± 14.9 | 18.2 ± 14 | P < 0.001 |

AST = antimicrobial susceptibility testing; ASP = antimicrobial stewardship program

Length of hospitalization (LOS) (10.7 ± 11.1 d vs. 15.5 ± 18.1 d, p=0.05) and length of in-patient antimicrobial therapy (6.7 ± 3.8 d vs. 8.8 ± 7.8 d, p=0.036) were significantly shorter for the ASP and MALDI-TOF/Vitek 2 group. Of interest, the LOS differences between the two groups was most dramatic in ICU, averaging 7 days shorter for patients in the ASP and MALDI-TOF/Vitek 2 group, (p<0.001) whereas patients admitted to the general medicine floor averaged 4 d shorter in the ASP and MALDI-TOF/Vitek 2 group. Finally, we sought to determine which variables were associated with LOS using a stepwise linear regression analysis. Variables that were significant by univariate analysis were entered into the linear regression analysis to determine variables that significantly correlated with LOS. After the stepwise procedure, one variable was included in the model, length of in-patient antimicrobial (Table 3). This significant variable captured 31% of the LOS variation (p<0.0001).

**Table 3.**
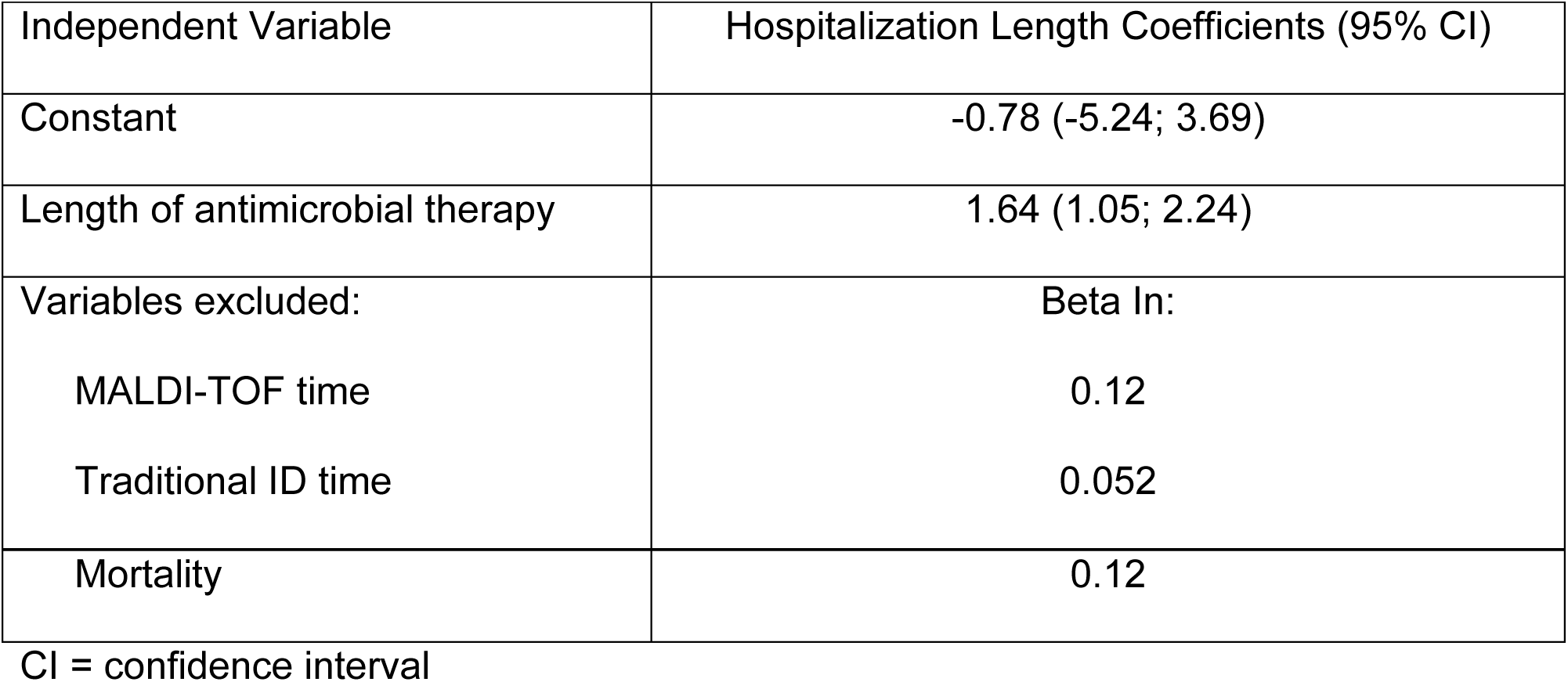
Length of Hospitalization Regression Analysis

## Discussion

The use of rapid diagnostic testing including MALDI-TOF and Vitek 2 AST has been proven to reduce time to organism identification. However, the incorporation of the ASP team along with rapid isolate identification and AST allows rapid interpretation of the results communicated to the primary care physician to make changes in antimicrobial therapy. Huang AM, et al. and Perez and colleagues both conducted pre-post quasi-experimental studies integrating MALDI-TOF organism identification plus ASP intervention, but limited to either BSIs, or Gram-negative bacteremias, respectively(6, 7). Huang, et al. enrolled 245 patients into an intervention group using MALDI-TOF and ASP and 256 patients into a preintervention group for bacteremias. In their study, length of ICU stay averaged 6 d shorter for the intervention group. Perez and colleagues enrolled 153 patients with antibiotic resistant Gram-negative bacteremias as the control group and 112 patients in the intervention group. Their LOS differences averaged 6 d longer for the control group overall with the ICU LOS averaging 5.3 d longer. These studies are consistent with the current study, which demonstrated that, with shorter turnaround time for ID and AST using optimized microbiology workflow and MADLI-TOF and Vitek 2, combined with a vigorous ASP, a significant reduction in LOS was achieved. Our previous de-escalation study confirms these results that shorter LOS is more exaggerated for ICU patients(12). Wenzler and colleagues reported use of MALDI-TOF and ASP for pneumonia or bacteremia caused by *A. baumannii.* Their results demonstrated the combination reduced time to effective therapy by 41 h and was associated with an increased clinical cure for those patients. Bookstaver et al. combined rapid diagnostic testing using Biofire BCID and ASP as a bundle for their quasi-experimental cohort of Gram-negative bloodstream infections in their multihospital health system(13). These investigators enrolled 830 patients into a preintervention group and 333 patients into a postintervention group with Gram-negative bacteremias. Their results demonstrated significant reductions in median time to de-escalate combination antimicrobials and initiation of more appropriate empiric antimicrobial therapy (before AST results were known). These investigators found significant reductions in median time to de-escalate combination antimicrobials and more appropriate empiric antimicrobial therapy (before AST results were known). Taken together along with the results of these studies, implementation of this process throughout our 14-hospital health-system with centralized Microbiology Laboratory would show a significant benefit to the care of infected patients now that there are more ASP pharmacists (3 covering CHI Health) and an ID physician with 0.5 FTE devoted to ASP across all hospitals in the health system.

However, there are limits associated with this study. A lack of patient randomization in the quasi-experimental study may not take into account changes in standard of care from the previous year, which could impact LOS. To our knowledge, both years incorporated Biofire BCID and there were no significant changes in standard of care except the introduction of the ASP team to make antimicrobial recommendations. The study periods were selected to minimize bias associated with medical residents and fellows at our academic medical center. This study excluded patients transferred into our regional hospital so all antimicrobials administration times were in-hospital times and not from administrations prior to admission. The effect of ASP was not directly identified in the stepwise multiple regression analysis. However, length of antimicrobials was identified as the primary factor associated with LOS, and ASP team did have an impact on this variable. By making appropriate antimicrobial recommendations, the ASP team impacted length of therapy and indirectly (probably) impacted LOS. It would be of interest to repeat this study from all CHI Health hospitals in Nebraska to determine the effect of ASP team and MALDI-TOF/Vitek 2 would have on these community hospitals in addition to this report since the centralized laboratory receives these specimens as far away as Kearney, NE (186 miles).

In conclusion, use of ASP and MALDI-TOF/Vitek 2 rapid identification and AST demonstrated for urine, blood, and sputum cultures a significant reduction in time to isolate identification and AST results, which translated to significant reduction in antibiotic length of therapy and hospital LOS.

